# Distinct roles of prefrontal cortex neurons in set shifting

**DOI:** 10.1101/2024.08.20.608808

**Authors:** Marco Nigro, Lucas Silva Tortorelli, Hongdian Yang

## Abstract

Cognitive flexibility, the ability to adjust behavioral strategies in response to changing environmental contingencies, requires adaptive processing of internal states and contextual cues to guide goal-oriented behavior, and is dependent on prefrontal cortex (PFC) functions. However, the neurophysiological underpinning of how the PFC supports cognitive flexibility is not well understood and has been under active investigation. We recorded spiking activity from single PFC neurons in mice performing the attentional set-shifting task, where mice learned to associate different contextually relevant sensory stimuli to reward. We identified subgroups of PFC neurons encoding task context, choice and trial outcome. Putative fast-spiking neurons were more involved in representing outcome and choice than putative regular-spiking neurons. Regression model further revealed that task context and trial outcome modulated the activity of choice-encoding neurons in rule-dependent and cell type-dependent manners. Together, our data provide new evidence to elucidate PFC’s role in cognitive flexibility, suggesting differential cell type-specific engagement during set shifting, and that both contextual rule representation and trial outcome monitoring underlie PFC’s unique capacity to support flexible behavioral switching.

## Introduction

The ability to adjust behavioral strategies in response to changing environmental contingencies, termed cognitive flexibility, serves as an essential executive function. Flexibility, or rule switching, requires adaptive processing of internal states and contextual cues to guide goal-oriented behavior, and is vital to the survival of organisms. Inappropriate behavioral adjustments, such as deficits in modifying responses to a rule change, are a hallmark of impaired executive functions observed in a broad spectrum of psychiatric disorders (Miller and Cohen, 2001; Uddin, 2021).

Considerable efforts have been made to uncover the neural substrates of flexible behavioral switching (see reviews (Mesulam, 1998; Miller, 1999; Miller and Cohen, 2001; Ragozzino, 2007; Le Merre et al., 2021; Uddin, 2021)). Set shifting, a type of rule switching that requires attending to or ignoring a stimulus feature in a context-dependent way, is widely used to assess cognitive flexibility. The Wisconsin Card Sorting Test (WCST), the Intra-Extra Dimensional Set Shift Task (IED) and their analogs have been implemented to test human and animal subjects (Berg, 1948; Milner, 1963; Roberts et al., 1988; Dias et al., 1996a; Monchi et al., 2001; Barnett et al., 2010; Brown and Tait, 2015). Decades of research have established that the prefrontal cortex (PFC) is required for set shifting (Berg, 1948; Milner, 1963; Dias et al., 1996a, 1996b; Ridderinkhof, 2004; Ragozzino, 2007; Floresco et al., 2009; Dajani et al., 2020). However, the neurophysiological underpinning of how the PFC mediates different aspects of flexible decision-making processes to support set shifting is not well understood. Importantly, although loss-of-function work has shown that the medial PFC (mPFC) is associated with attentional switching across, but not within perceptual dimensions (e.g., (Owen et al., 1991; Dias et al., 1996b, 1997; Birrell and Brown, 2000; Ridderinkhof, 2004; Ragozzino, 2007; Bissonette et al., 2008)), the neural substrates that support such functional specificity remain elusive and are under active investigation (e.g., (Cho et al., 2020, 2023; Benoit et al., 2022)).

In an effort to advance our understanding of PFC’s role in flexible behavior, we trained mice to perform the attentional set-shifting task (AST), which follows the principles of WCST and IED, and trains animals to continuously adapt to multiple rule changes (Birrell and Brown, 2000; Colacicco et al., 2002; Garner et al., 2006; Bissonette et al., 2008; Lapiz-Bluhm et al., 2008; Heisler et al., 2015) (Fig. 1A). These rule changes may or may not involve the mPFC (Birrell and Brown, 2000; McAlonan and Brown, 2003; Bissonette et al., 2008, 2013). Specifically, in extra-dimensional shift (EDS) subjects learn to attend to a novel stimulus from a different dimension (e.g., from digging medium to odor) to seek reward, and task performance is impaired by mPFC lesion. In contrast, intra-dimensional reversal (REV) requires attending to a previously unrewarding stimulus and ignoring a previously rewarding stimulus within the same stimulus dimension, and is not affected by mPFC lesion.

**Figure 1.**
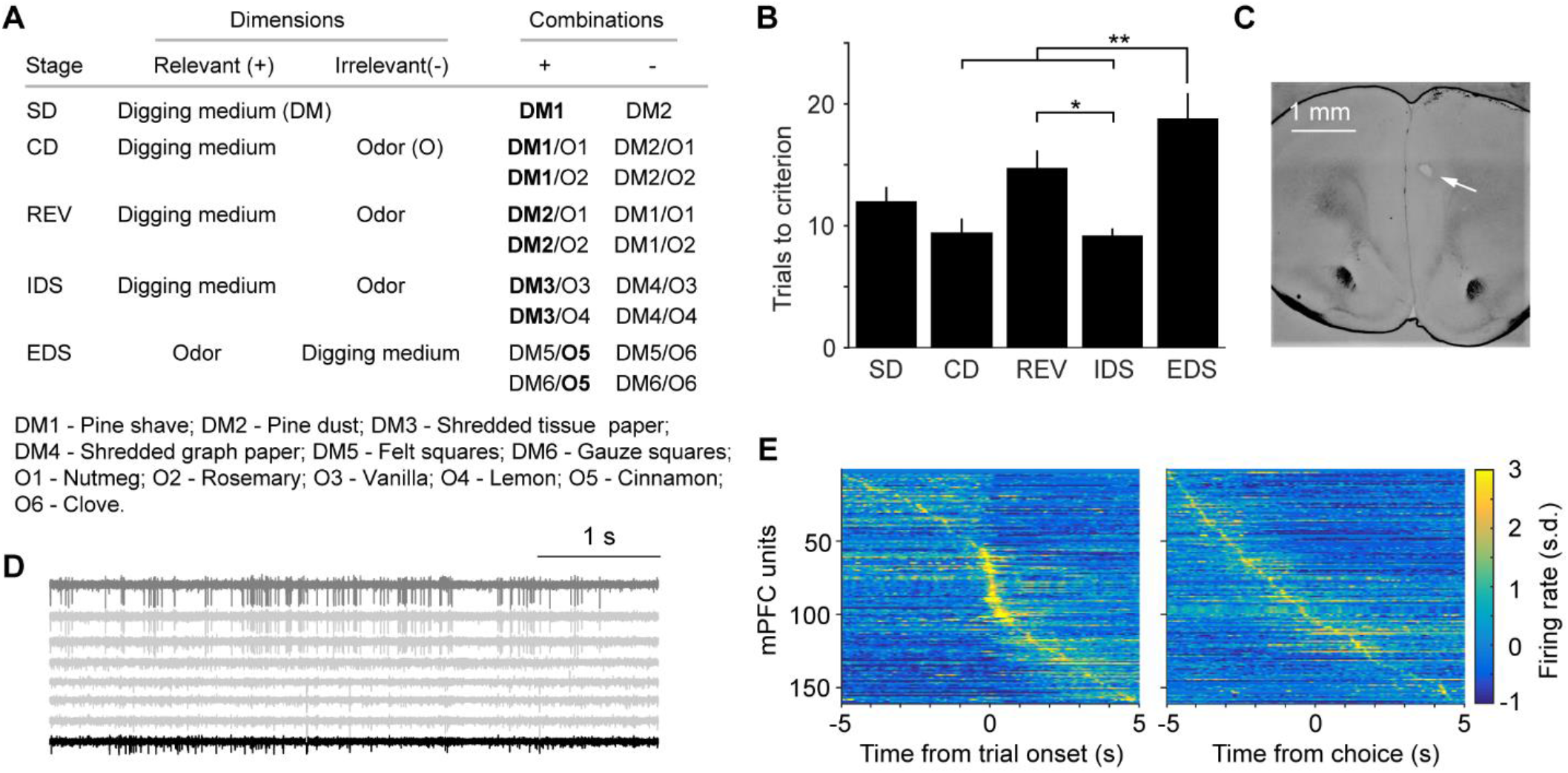
Tetrode recording in the mPFC during AST. (A) Test structure of AST. (B) Task performance (total number of trials to criterion) varied across stages. Repeated-measure ANOVA, F(4, 60) = 8.6, p = 1.5e-5, n = 15. Post hoc Tukey-Kramer tests revealed that mice took more trials to complete REV and EDS stages. REV vs. IDS, p = 0.018; EDS vs. CD, p = 0.0038; EDS vs. IDS, p = 0.0052. All other paired comparisons were not significantly different. (C) Coronal brain section showing an electrolytic lesion marking the recording site (arrow) in the prelimbic region. (D) Eight example traces from a 32-channel tetrode recording in the mPFC during behavior. (E) Example heat map of trial-averaged spiking activity (z-scored) of all 161 units during trial onset (left) and during correct choice (right, ± 5 s) in EDS.

We recorded spiking activity from single units in mice performing AST. We identified subgroups of mPFC neurons representing different task-related variables, namely task context, trial outcome and choice. We found that putative fast-spiking neurons were more engaged in representing outcome and choice than putative regular-spiking neurons. We showed that both context and outcome signals significantly modulated the activity of choice-encoding neurons in EDS. The modulatory effects were most obvious in fast-spiking neurons and were absent in REV. Together, our data suggest differential cell type-specific engagement during rule switching, and that both contextual rule representation and outcome monitoring underlie mPFC’s unique role in supporting set-shifting behavior.

## Results

We trained mice to perform the attentional set-shifting task (AST) using procedures similar to previous work (Methods. Liston et al., 2006; Snyder et al., 2012). Briefly, in most stages of the task, mice learned to associate one relevant sensory stimulus out of several possible ones to reward (Fig. 1A, Fig. S1). The relevant stimulus remained in the dimension of digging medium in early stages of the task (simple discrimination, SD; compound discrimination, CD; intra-dimensional reversal, REV; intra-dimensional shift, IDS), and shifted to the dimension of odor in the last stage of extra-dimensional shift (EDS). Mice promptly learned to follow the rule in each stage. However, REV and EDS appeared to be more challenging as mice needed more trials to reach performance criterion (six consecutive correct trials, Fig. 1B) (Birrell and Brown, 2000; Liston et al., 2006; Snyder et al., 2012). To elucidate the role of mPFC in cognitive flexibility, we conducted tetrode recording during task performance (161 single units from 15 sessions, Fig. 1C-E, Methods). Previous loss-of-function studies have reported that the mPFC is specifically required for the successful completion of EDS (e.g., (Dias et al., 1996b; Birrell and Brown, 2000; Bissonette et al., 2008)), and our analyses were focused on EDS to assess the neural substrates.

First, we sought out to examine the extent to which abstract contextual rule information was represented in the mPFC. In AST, this refers to the stimulus dimension that subjects learn to attend to (digging medium vs. odor). Plateaued performance following a rule change has been taken as important evidence that subjects readily adapt to the new rule (e.g., (Mansouri et al., 2006; Sleezer et al., 2016)). Indeed, we found that the spiking activity of a subset of mPFC neurons tracked the attended stimulus dimension when performance was plateaued (last set of consecutive correct trials, Fig. 2A, B). Using Receiver-Operating-Characteristic (ROC) analysis (Green and Swets, 1966), we identified 31% (50/161) of mPFC neurons whose activity was significantly correlated with task context (Fig. 2C-G, Methods), and we referred to them as context neurons. Similar numbers of neurons exhibited higher or lower activity when the relevant stimulus dimension shifted from digging medium to odor (context+ vs. context-, 27 vs. 23 neurons). We did not include SD in the analysis because the odor dimension was not explicitly introduced (Fig. 1A, Methods). However, the identified context neurons exhibited comparable activity in SD as in other digging medium-relevant stages (Fig. S2), supporting their robust representation of stimulus dimension. Further, the classification of context neurons was supported by a generalized linear model (GLM), where the coefficients of stimulus dimension were significantly stronger than other task-related variables, and stronger than those of non-context neurons (Fig. S3, Methods). We trained a decoder to evaluate the extent to which we can predict the shift of task context based on context neuron activity, and the decoder was able to achieve 80.8 ± 5.9% accuracy (Fig. 2H, I, Methods). Context-related activity sustained for tens of seconds before explicit task engagement (Fig. S4), suggesting that context information was represented in persistent activity, in support of other studies (e.g., (Mansouri et al., 2006; Sleezer et al., 2016; Bari et al., 2019)). We next evaluated context neuron activity during rule learning and found that their activity exhibited gradual changes when the relevant dimension shifted from digging medium to odor (from IDS to EDS, Fig. 2J, K). Since the dimensional rule shift was not cued, this finding supports that context representation develops over learning.

**Figure 2.**
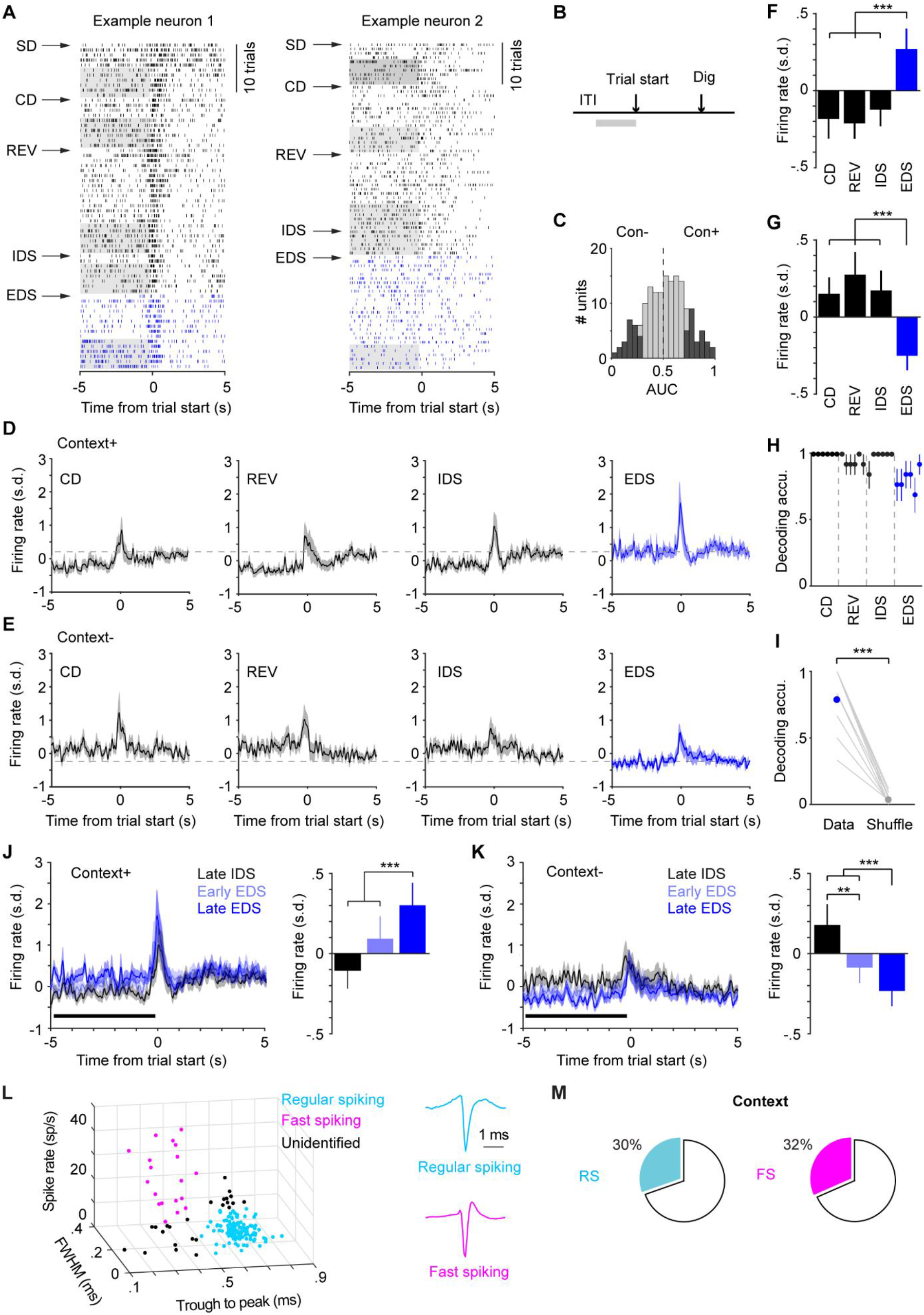
Task context encoding in the mPFC. (A) Spike rasters from two example neurons showing enhanced (left) or suppressed (right) activity during intertrial intervals in the last 6 consecutive correct trials (grey area) in EDS compared with other stages. Ticks represent spikes. (B) Illustration of the time window used to classify context encoding. (C) Distribution of AUC values of context encoding for all neurons (light grey). Significantly modulated neurons (p < 0.05) were in dark. (D) Group mean peri-event spike time histogram (PETH) of context+ neurons (n = 27) aligned to trial onset in stages CD through EDS. Mean firing rate during a 5-s window before trial start (black horizontal bar) is shown in E. Dashed line is to aid comparison. (E) Group mean peri-event spike time histogram (PETH) of context-neurons (n = 23) aligned to trial onset in stages CD through EDS. Mean firing rate during a 5-s window before trial start (black horizontal bar) is shown in G. (F) Context+ neurons showed significantly higher activity in EDS. Repeated-measures ANOVA, F(3, 78) = 16.1, p = 3.03e-8, n = 27. Post hoc Tukey-Kramer tests: EDS vs. CD, p = 2.1e-4; EDS vs. REV, p = 2.1e-6; EDS vs. IDS, p = 2.2e-4. All other paired tests were not significant. (G) Context-neurons showed significantly lower activity in EDS. Repeated-measures ANOVA, F(3, 66) = 15.14, p = 1.3e-7, n = 23. Post hoc Tukey-Kramer tests: EDS vs. CD, p = 6.4e-6; EDS vs. REV, p = 8.9e-6; EDS vs. IDS, p = 6.1e-5. All other paired tests were not significant. (H) Decoding of task context of the last six trials in stages CD through EDS (n = 15). (I) Average decoding accuracy of last six trials in EDS for each recording (n = 15), compared with shuffled model (Data vs. Shuffle, 80.8 ± 5.9% vs. 2.6 ± 0.9%, p = 2.4e-4). (J) Left: Group mean PETH of context+ neurons aligned to trial onset from late IDS (black, last 6 correct trials), early EDS (light blue, all trials preceding last 6 correct trials), and late EDS (last six correct trials). Right: Mean firing rate during a 5-s window before trial start. Repeated-measures ANOVA, F(2, 52) = 13.1, p = 2.5e-5, n = 27. Post hoc Tukey-Kramer tests: Late IDS vs. Early EDS, p = 0.14; Late IDS vs. Late EDS, p = 1.1e-4; Early EDS vs. Late EDS, p = 2.4e-4. (K) Left: Group mean PETH of context-neurons aligned to trial onset from late IDS (black, last 6 correct trials), early EDS (light blue, all trials preceding last 6 correct trials), and late EDS (last six correct trials). Right: Mean firing rate during a 5-s window before trial start. Repeated-measures ANOVA, F(2, 44) = 20.9, p = 4.1e-7, n = 23. Post hoc Tukey-Kramer tests: Late IDS vs. Early EDS, p = 0.0095; Late IDS vs. Late EDS, p = 3.2e-5; Early EDS vs. Late EDS, p = 1.4e-5. (L) Classifying putative fast-spiking (magenta) and regular-spiking (cyan) neurons based on spike waveform features and spike rate. (M) Similar proportions of RS and FS neurons encoded context. 34 out of 112 RS vs. 6 out of 19 FS, 30% vs. 32%, p = 0.91.

To examine the contributions of different cell types to set shifting, we classified the recorded units into putative inhibitory fast spiking (FS) and putative excitatory regular spiking (RS) based on spike waveform features and firing rate (Barthó et al., 2004; Ji and Neugebauer, 2012): FS, trough to peak = 0.35 ± 0.02 ms; baseline firing rate = 28.14 ± 2.77 spikes/s, n = 19; RS, trough to peak = 0.67 ± 0.01 ms; baseline firing rate = 2.85 ± 0.24 spikes/s, n = 112 (Methods, Fig. 2L). The remaining units were considered unidentified and excluded from cell type-related analyses. We found similar proportions of RS and FS neurons encoding task context (34/112 RS vs. 6/19 FS, 30% vs. 32%, p = 0.91, chi-squared test, Fig. 2M).

Next, we evaluated to what extent mPFC activity represented previous trial outcome. Using similar ROC analysis, we identified 22% (36/161) of neurons exhibiting differential activity following correct (rewarded) or incorrect (unrewarded) trials (Fig. 3A, B, Methods). 64% of these neurons (23/36) showed higher activity when previous trials were correct compared with when previous trials were incorrect (outcome+, Fig. 3C). The remaining 36% of outcome neurons (13/36) exhibited the opposite trend, increasing firing rate following incorrect trials compared to following correct trials (outcome-, Fig. 3D). Based on outcome neuron activity, a decoder was able to predict trial outcome with 83.0 ± 3.4% accuracy (Fig. 3E, Methods). Similar to context encoding, outcome-related activity sustained for tens of seconds prior to task engagement (Fig. S5), indicating that outcome information (in particular negative outcome) was represented in persistent mPFC activity. 28% (10/36) of outcome-encoding neurons also represented context, supporting mixed tuning in the PFC (Rigotti et al., 2013; Fusi et al., 2016; Tye et al., 2024). Interestingly, higher proportions of FS neurons were found to represent outcome (26/112 RS vs. 9/19 FS, 23% vs. 47%, p = 0.028, Fig. 3F).

**Figure 3.**
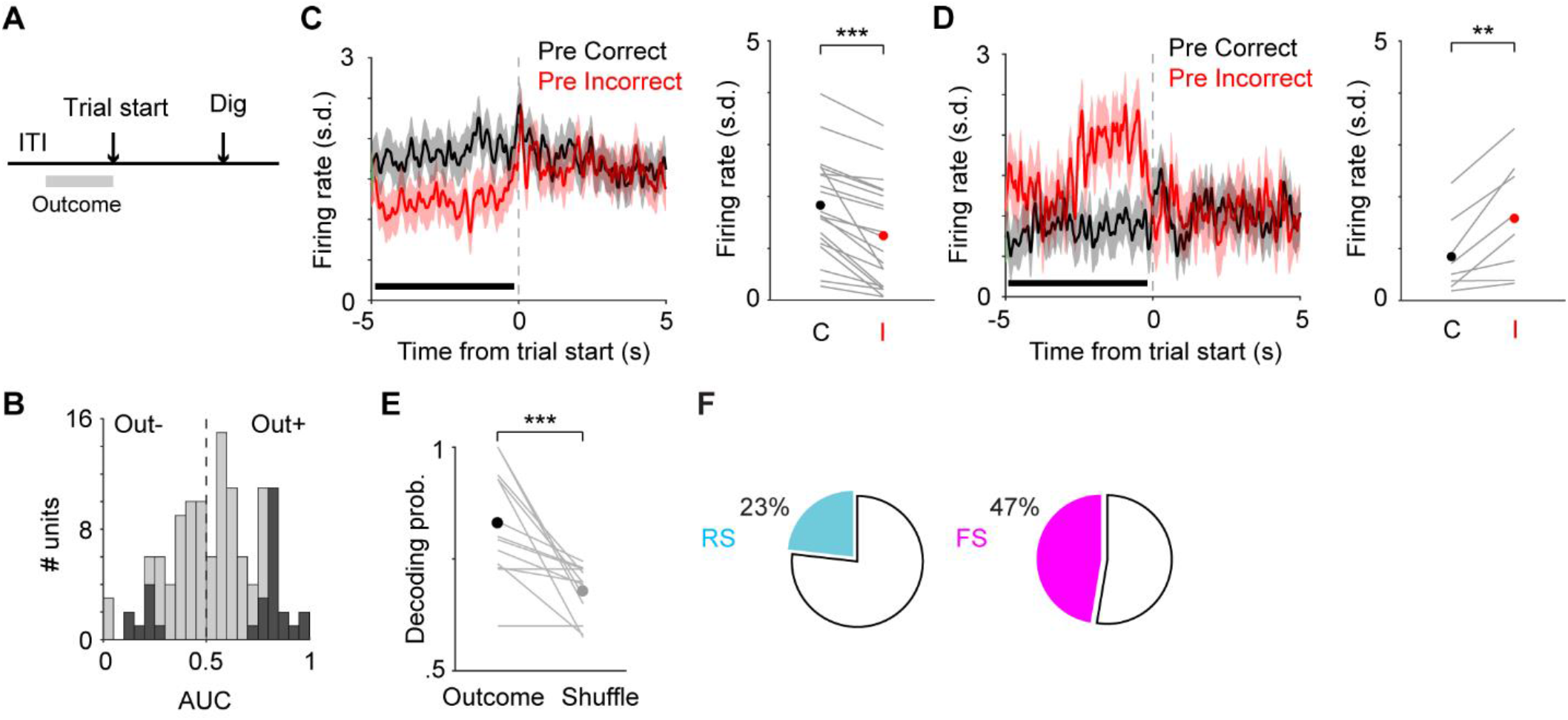
Trial outcome encoding in the mPFC. (A) Illustration of the time window used to classify outcome encoding. (B) Distribution of AUC values of outcome encoding for all neurons (light grey). Significantly modulated neurons (p < 0.05) were in dark. (C) Left: Group mean PETH of outcome+ neurons in EDS (n = 23) aligned to trial onset when previous trials were correct (black) and incorrect (red). Right: Mean firing rate during a 5-s window before trial start when previous trials were correct (black) and incorrect (red). p = 2.7e-5. Lines: individual neurons. Dots: mean. (D) Left: Group mean PETH of outcome-neurons in EDS (n = 13) aligned to trial onset when previous trials were correct (black) and incorrect (red). Right: Mean firing rate during a 5-s window before trial start when previous trials were correct (black) and incorrect (red). p = 2.4e-4. Lines: individual neurons. Dots: mean. (E) Average outcome decoding accuracy of EDS for each recording (n = 15), compared with shuffled model (Outcome vs. Shuffle, 83.0 ± 3.4% vs. 66.9 ± 1.7%, p = 4.9e-4). (F) Higher proportions of FS neurons encoded outcome. 26 out of 112 RS vs. 9 out of 19 FS, 23% vs. 47%, p = 0.028.

We then identified choice neurons, whose activity correlated with the upcoming choices on current trials (correct vs. incorrect, 23/161, Fig. 4A, B). 56% of these neurons (13/23) exhibited higher activity preceding correct choices than incorrect choices (Fig. 4C). These neurons, hereafter referred to as choice+, also showed significantly elevated activity immediately before correct choices compared to after these choices. In contrast, their activity before and after incorrect choices were similar (Fig. S6A). Choice-neurons (10/23) exhibited lower activity preceding correct choices than incorrect choices (Fig. 4D), and did not show any differential activity before and after correct or incorrect choices (Fig. S6B). A decoder was able to predict trial-by-trial choices with 80.8 ± 2.1% accuracy from choice neuron activity (Fig. 4E, Methods). We also found higher proportion of FS neurons encoding choice (12/112 RS vs. 6/19 FS, 11% vs. 32%, p = 0.015, Fig. 4F). A considerable fraction of choice-encoding neurons also represented other task-related variables (43% (10/23) choice neurons represented outcome, and 39% (9/23) choice neurons also represented context), in further support of mixed tuning. Together, our results showed that putative FS neurons were more involved in representing outcome and choice during set shifting.

**Figure 4.**
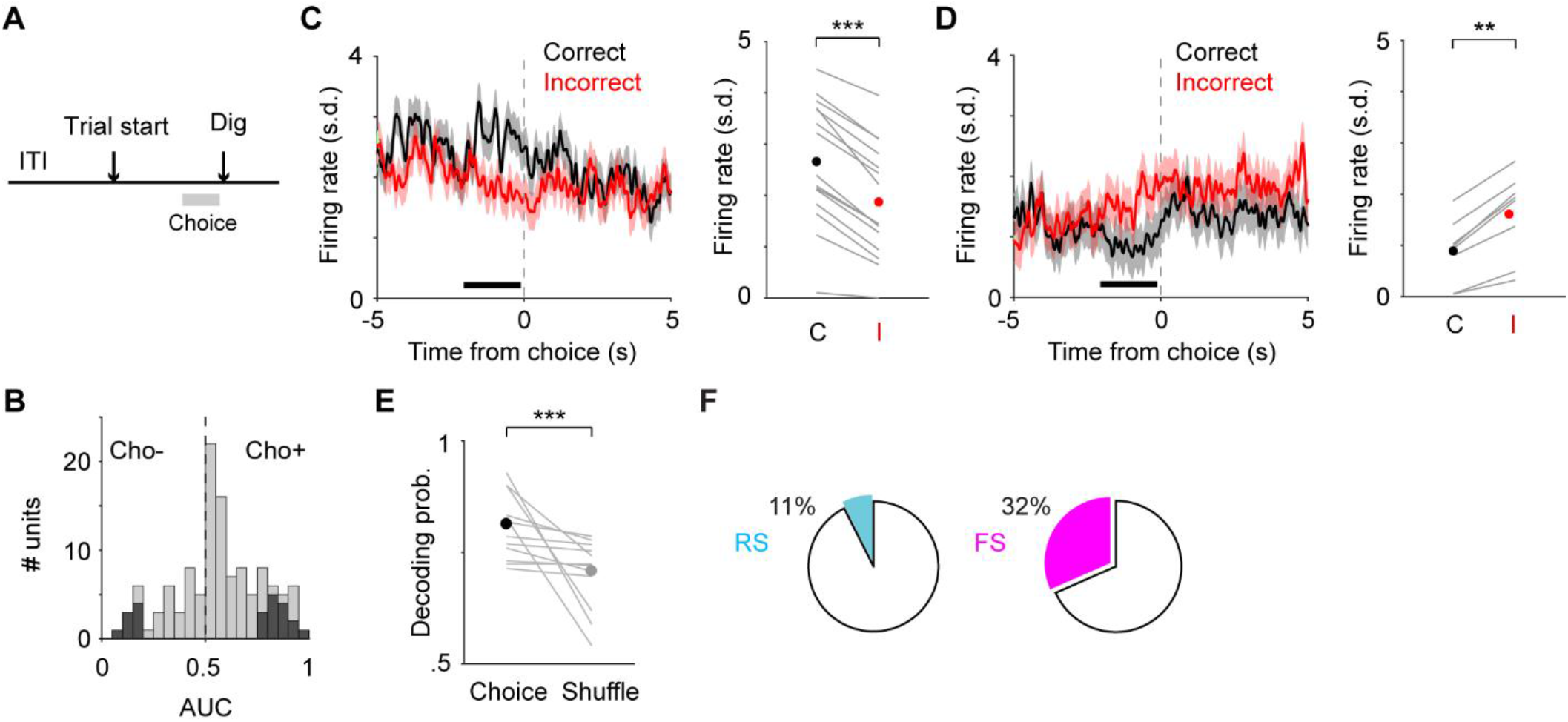
Choice encoding in the mPFC. (A) Illustration of the time window used to classify choice encoding. (B) Distribution of AUC values of choice encoding for all neurons (light grey). Significantly modulated neurons (p < 0.05) were in dark. (C) Left: Group mean PETH of choice+ neurons in EDS (n = 13) aligned to trial onset when the upcoming choices of current trials were correct (black) and incorrect (red). Right: Mean firing rate during a 2-s window before digging when the upcoming choices of current trials were correct (black) and incorrect (red). p = 2.4e-4. Lines: individual neurons. Dots: mean. (D) Left: Group mean PETH of choice-neurons in EDS (n = 10) aligned to trial onset when the upcoming choices of current trials were correct (black) and incorrect (red). Right: Mean firing rate during a 2-s window before digging when the upcoming choices of current trials were correct (black) and incorrect (red). p = 0.002. Lines: individual neurons. Dots: mean. (E) Average choice decoding accuracy of EDS for each recording (n = 15), compared with shuffled model (Choice vs. Shuffle, 80.8 ± 2.1% vs. 70.6 ± 2.3%, p = 9.8e-4). (F) Higher proportions of FS neurons encoded choice. 12 out of 112 RS vs. 6 out of 19 FS, 11% vs. 32%, p = 0.015.

To understand how context and outcome may affect decision making, we examined the impact of these two variables on the activity of choice neurons. We divided each trial into four 2-s bins, with two bins prior to trials start (T1, T2), and two other bins prior to choice (T3, T4. Fig. 5A). We used GLM to calculated the regression coefficients for the regressors of trial outcome and contextual rule on choice neurons in EDS. GLM confirmed that the identified choice+ neurons prominently represented the choice signal prior to digging (Fig. S7). Interestingly, these neurons showed non-zero coefficients for outcome and context. Specifically, we found significant coefficients for outcome before trial start (T1, T2) and before choice (T4, Fig. 5B). For context, we observed significant non-zero coefficients before trial start (T2) and before choice (T3, T4, Fig. 5C). These effects were mostly absent in choice-neurons (Fig. S8). Albeit the small sample sizes, the modulatory effects were present in FS neurons, as choice+ FS neurons exhibited significant coefficients for outcome (T1-T4) and context (T2, Fig. 5D, E). In contrast, choice-encoding RS neurons were not modulated by outcome or context (Fig. 5F, G). In summary, our findings revealed distinct modulation patterns in putative FS and RS neurons, with context and outcome-related information primarily affecting FS activity.

**Figure 5.**
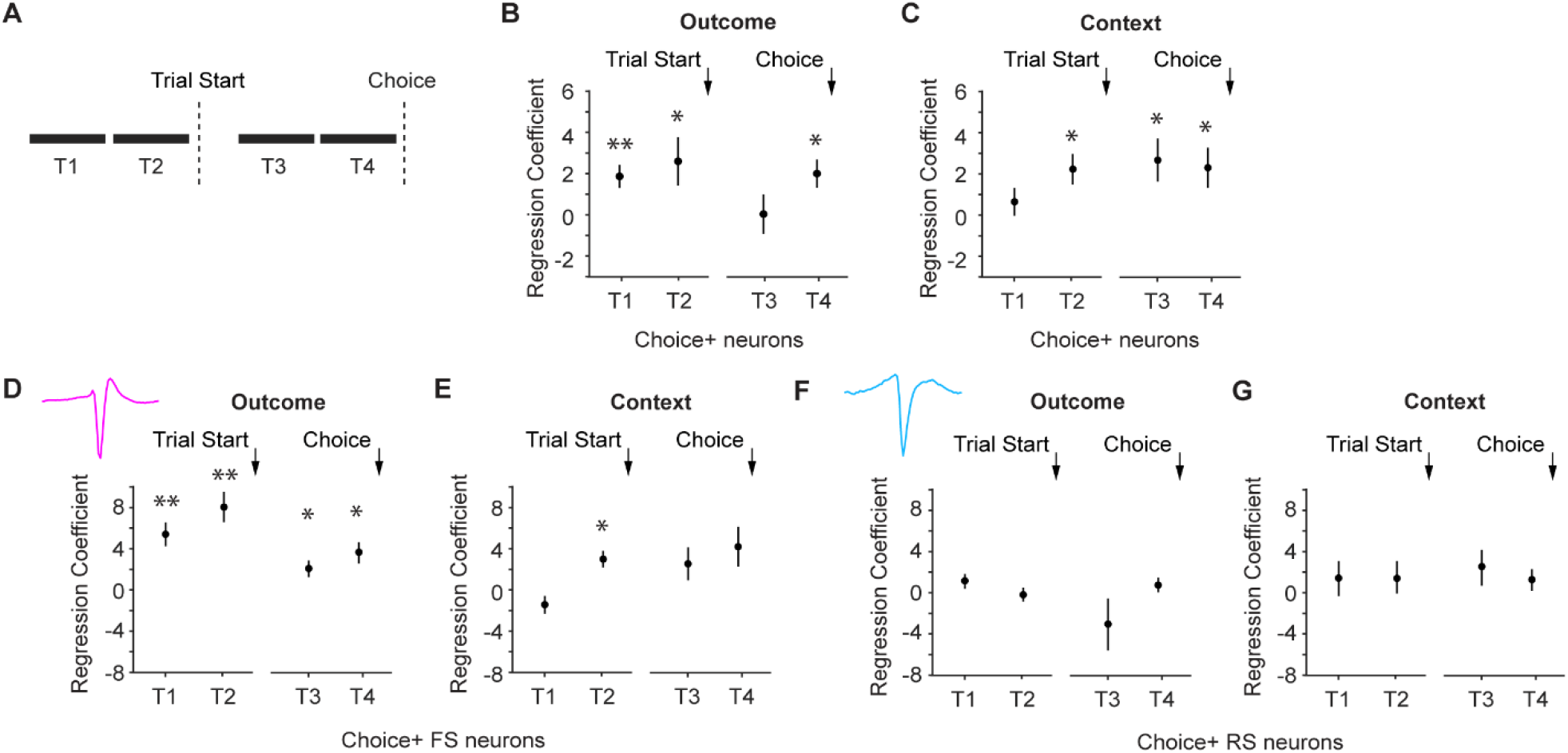
Context and outcome modulate choice-encoding neuronal activity in EDS. (A) Illustration of the four trial epochs. T1: -4 to -2 s from trial onset; T2: -2 to 0 s from trial onset; T3: -4 to -2 s from digging, T4: -2 to 0 s from digging; (B) Regression coefficients of the outcome regressor for choice+ neurons (n = 13). Coefficients in T1, T2 and T4 were significantly different from 0. T1, p = 0.01; T2, p = 0.05; T3, p = 0.97; T4, p = 0.01. (C) Regression coefficients of the context regressor for choice+ neurons (n = 13). Coefficients in T2, T3 and T4 were significantly different from 0. T1, p = 0.48; T2, p = 0.02; T3, p = 0.03; T4, p = 0.04. (D) Regression coefficients of the outcome regressor for fast-spiking choice+ neurons in EDS (n = 5). Coefficients were significantly different from 0 in all epochs. T1, p = 0.009; T2, p = 0.005; T3, p = 0.038; T4, p = 0.025. (E) Regression coefficients of the context regressor for fast-spiking choice+ neurons in EDS (n = 5). Coefficients in T2 were significantly different from 0. T1, p = 0.12; T2, p = 0.02; T3, p = 0.17; T4, p = 0.1. (F) Regression coefficients of the outcome regressor for regular-spiking choice+ neurons (n = 5). Coefficients were not significantly different from 0 in any epochs. T1, p = 0.20; T2, p = 0.49; T3, p = 0.23; T4, p = 0.5. (G) Regression coefficients of the context regressor for regular-spiking choice+ neurons in EDS (n = 5). Coefficients were not significantly different from 0 in any epochs. T1, p = 0.49; T2, p = 0.44; T3, p = 0.26; T4, p = 0.34. T test for all comparisons in Fig. 5.

Lastly, we wondered whether the observed modulation patterns were specific to EDS switching. We analyzed REV as a comparison, which was also behaviorally demanding but not affected by mPFC perturbation (Fig. 1B, Birrell and Brown, 2000; Bissonette et al., 2008). We identified largely distinct groups of neurons encoding outcome and choice in REV (Outcome, REV vs EDS: 22 vs. 36 neurons, 4 overlapped neurons; Choice, REV vs EDS: 19 vs. 23 neurons, 4 overlapped neurons). More mPFC neurons encoded outcome in EDS than REV (Outcome, REV vs. EDS, 14% (22/161) vs. 22% (36/161), p = 0.04; Choice, REV vs. EDS, 12% (19/161) vs. 14% (23/161), p = 0.51). Regression analysis revealed that trial outcome did not significantly affect the activity of choice neurons in REV (Fig. 6A, Fig. S9). Since REV did not involve a change of stimulus dimension, we treated the result that task context did not affect choice neuron activity in REV as a positive control (Fig. 6B, Fig. S9). Finally, we assessed how different cell types were engaged in REV and EDS. For choice, we found similar proportions of RS neurons in REV and EDS (REV vs. EDS, 15/112 RS vs. 12/112 RS, 13% vs. 11%, p = 0.54). However, REV engaged a lower proportion of FS choice-encoding neurons (REV vs. EDS, 1/19 FS vs. 6/19 FS, 5% vs. 32%, p = 0.037, Fig. 6C). Similarly, lower proportions of FS outcome-encoding neurons were identified in REV (REV vs. EDS, 15/112 RS vs. 26/112 RS, 13% vs. 23%, p = 0.057; 0/19 FS vs. 9/19 FS, 0% vs. 47%, p = 5.9e-4, Fig. 6D). Together, our data uncovered substantial differences in mPFC representation during different types of rule switching behavior, such that task context and trial outcome modulated the activity of choice-encoding neurons only in EDS but not REV, and primarily affected FS but not RS neurons.

**Figure 6.**
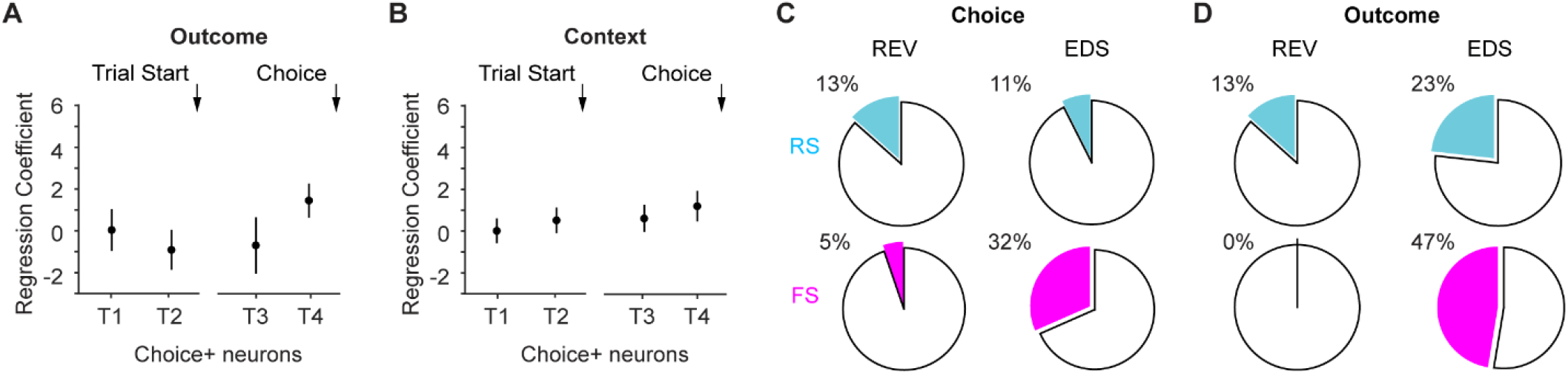
Context and outcome do not modulate choice-encoding neuronal activity in REV. (A) Regression coefficients of the outcome factor for choice+ neurons in REV (n = 13). Coefficients were not significantly different from 0 in any epochs. T1, p = 0.99; T2, p = 0.35; T3, p = 0.63; T4, p = 0.09. (B) Regression coefficients of the context factor for choice+ neurons in REV (n = 13). Coefficients were not significantly different from 0 in any epochs. T1, p = 0.92; T2, p = 0.38; T3, p = 0.3; T4, p = 0.1. (C) Comparison of the proportions of cell type-specific choice-encoding neurons in REV and EDS. (D) Comparison of the proportions of cell type-specific outcome-encoding neurons in REV and EDS.

## Discussion

To elucidate mPFC’s role in cognitive flexibility, we recorded spiking activity from single units in mice performing AST. We identified neuronal subgroups encoding different task-related variables, namely task context, trial outcome and choice. Importantly, we showed that putative FS interneurons were more engaged than putative RS neurons in representing outcome and choice.

By contrasting neuronal responses in EDS to REV, regression model revealed that context and outcome signals modulated the activity of choice-encoding neurons in task-dependent and cell type-dependent manners. Together, our data suggest differential cell type-specific engagement during flexible rule switching, and that both contextual rule representation and trial outcome monitoring underlie mPFC’s unique capacity to support set shifting.

mPFC has been proposed to support cognitive flexibility by encoding abstract contextual rules (Wallis et al., 2001; Meyers et al., 2008; Rich and Shapiro, 2009; Durstewitz et al., 2010; Hyman et al., 2012; Mante et al., 2013; Rodgers and DeWeese, 2014; Siniscalchi et al., 2016; Rikhye et al., 2018; Reinert et al., 2021), or by encoding feedback signals (Luk and Wallis, 2009; Bissonette and Roesch, 2015; Del Arco et al., 2017; Bari et al., 2019; Norman et al., 2021; Spellman et al., 2021). These two hypotheses are not necessarily exclusive becasue when subjects are unaware of the rule change, they likely utilize more than one stream of information to solve the task (Ridderinkhof, 2004; Rushworth and Behrens, 2008; Mansouri et al., 2009; Bissonette et al., 2013; Uddin, 2021). Indeed, our data show that abstract contextual rule-related information and trial outcome-related information are both represented in persistent activity in the mPFC. It is possible that using novel rather than familiar cues in EDS is important for the formation and utility of stimulus dimension in the mPFC (Birrell and Brown, 2000; Bissonette et al., 2013).

Our data further suggest the functional specificity of such representations, as context and outcome affected the activity of choice-encoding neurons only in EDS but not REV. Behaviorally, both REV and EDS appear to be more challenging as subjects typically take more trials to reach performance criterion (Birrell and Brown, 2000; Liston et al., 2006; Snyder et al., 2012). However, these two rule changes are thought to involve different cognitive processes as the former is referred to as affective shifting while the later as attentional shifting (Dias et al., 1996b; Floresco et al., 2009; Young et al., 2010). In REV, subjects are challenged to ignore the relevant stimulus from the previous stage, and to attend to a previously ignored stimulus within the same stimulus dimension. In EDS, subjects learn to direct their responses to a novel cue from the previously irrelevant stimulus dimension. According to learning theories, the improved performance in IDS (fewer trials to complete) strongly suggests that mice attend to the stimulus dimensions (digging medium vs. odor), and that solving EDS involves a shift in the attended dimension, rather than purely responding to specific sensory cues (Mackintosh, 1975; Roberts et al., 1988). Notably, the activity of choice-encoding neurons is modulated by context and outcome only in EDS but not REV, suggesting the unique neural substrates underlying mPFC’s functional specificity.

Why did context and outcome only affect choice+ neuron activity? The plateaued performance toward the end of a behavioral session was considered as rule acquisition, while earlier trials were considered as trial-and-error learning (Sleezer et al., 2016, 2017; Nigro et al., 2023). Thus, incorrect choices likely reflect the early rule learning phase, and correct choices likely reflect the late rule acquisition phase. We speculate that the increase in choice+ neuron activity prior to correct choices is therefore correlated with state changes in switching behavior, suggesting that outcome and context signals are important for driving rule switching in EDS.

Our findings further suggest a critical role for fast-spiking interneurons in set shifting, consistent with prior working demonstrating the importance of PV-mediated synchrony and differential encoding between RS and FS neurons in flexible behavior (Rikhye et al., 2018; Cho et al., 2020, 2023; Benoit et al., 2022). The stronger involvement of putative FS neurons implies a key role of inhibitory signaling in shaping information flow and excitation-inhibition balance, important in many neuropsychiatric conditions (e.g., (Rubenstein and Merzenich, 2003; Cho et al., 2015; Canetta et al., 2016; Cardin, 2018; Sohal and Rubenstein, 2019)).

One limitation of the current study is the relatively low number of simultaneously recorded neurons per behavioral session, which precludes performing comprehensive population-based analysis to better examine network dynamics (as in (Durstewitz et al., 2010; Jercog et al., 2021; Zhou et al., 2021; Richman et al., 2023)). Another limitation is that some cell type-related findings are based on a relatively low number of FS neurons. These limitations can be aided by recording from genetically identified neurons (e.g., (Pi et al., 2013; Pinto and Dan, 2015; Kim et al., 2016)) in future studies. Nevertheless, our single-cell analysis has uncovered new information on how individual neurons encode information during set shifting, elucidating the fundamental building blocks of neuronal computation and information processing.

Our work contributes to the growing interest in revealing neural mechanisms underlying more natural, ethologically relevant behavior (Parker et al., 2020; Dennis et al., 2021). Admittedly, such behavioral paradigms may not afford the level of task control more commonly seen in restrained, operant paradigms. Nevertheless, the challenge of dissociating movement-related signal from sensory- or decision-related signal is present in not only freely-moving, but also restrained settings (Musall et al., 2019; Steinmetz et al., 2019; Stringer et al., 2019; Zagha et al., 2022). Comprehensive behavioral tracking and motif analysis (e.g., (Wiltschko et al., 2015; Markowitz et al., 2023)) will help to identify whether specific behavioral patterns are associated with rule switching behavior. Ultimately, cognitive processes are not independent from sensory or motor processes. Cognition, perception and action may be implemented in a distributed rather than isolated manner (Cisek and Kalaska, 2010; Parker et al., 2020; Zagha et al., 2022).

## Author contributions

MN, LST and HY planned the project. LST performed experiments. MN and HY analyzed data and wrote the manuscript with assistance from LST.

## Acknowledgements

We thank Shaorong Ma for helping with the behavioral paradigm; Laurie Graham for instrument fabrication. HY was supported by UCR startup, Klingenstein-Simons Fellowship Awards in Neuroscience, and NIH grants (R01NS107355, R01NS112200).

## Materials and Methods

All procedures were performed in accordance with protocols approved by UC Riverside Animal Care and Use Committee (#20190031). Ten C57BL/6 mice of 8-12 weeks and mixed sex were used in this study. Procedures for microdrive construction and recording were similar to our previous work (Megemont et al., 2022, 2024). Briefly, the implants were custom microdrives with eight tetrodes, each consisting of four nichrome wires (200–300 kΩ). The microdrive was implanted through a ∼1 mm diameter craniotomy targeting the left mPFC (prelimbic area, 1.9-2.2 mm rostrocaudal and 0-0.5 mm mediolateral relative to bregma and 1 mm dorsoventral relative to brain surface). The microdrive was advanced in steps of 100 µm each day until reaching the recording depth of 1.4-1.6 mm. At the end of the experiment, an electrolytic lesion (100 μA, 20 s) was made prior to transcardial perfusion. Perfusions were done first with PBS followed by 4% PFA. The brain was sliced at 100 μm coronal sections to confirm the recording site.

Mice were singly housed after tetrode implant and allowed 2-3 days of recovery. Mice were then food restricted (80-85% of initial weight) and handled by the experimenter for 5-7 days. Next, mice were acclimated to the behavioral box (22 × 33 cm) and experimental setup for 1-2 days, followed by a brief training session to stimulate the innate burrowing/digging behavior to retrieve food reward from the ramekins. Two ramekins were placed at two corners of the behavioral box, both containing 25 mg of cheerios. Throughout the training session the reward was gradually buried in clean home cage bedding. In each trial mice were allowed 3-4 minutes to explore. Mice were considered well trained once they can consistently dig and retrieve the reward from both locations for 15-20 trials.

To assess flexible decision-making in freely moving mice, we adopted the 5-stage testing paradigm of the attentional set-shifting task (Liston et al., 2006; Snyder et al., 2012), consisting of the following stages: 1) simple discrimination (SD), in which animals choose between two digging medium associated with distinct textures (first stimulus dimension), only one of the two stimuli predicts food reward; 2) compound discrimination (CD), in which two odor cues (second stimulus dimension) are explicitly introduced. Each odor cue is randomly paired with a digging medium in every trial, but the reward is still predicted as in SD; 3) intra-dimensional reversal (REV), which preserves the task-relevant dimension (digging medium) but swaps cue contingencies; 4) intra-dimensional shift (IDS), which preserves the task-relevant dimension (digging medium), but replaces all four cues with novel ones (a new digging medium predicts reward); 5) extra-dimensional shift (EDS), which swaps the previous task-relevant and task-irrelevant dimensions with all cues replaced (a new odor cue predicts reward). All stages were performed within a single day, lasting 3-4 hours. In each trial, the ramekin associated with the relevant stimulus contained a retrievable reward. To avoid the possibility that mice used food odor cues to solve the task, the other ramekin contained a non-retrievable reward (trapped under a mesh wire at the bottom). The two ramekins were placed randomly in the two corners every trial. Between trials, mice were confined to the other side of the behavioral box (opposite to the ramekins) with a divider inserted (‘waiting zone’, Fig. S1), and had free access to water. Each trial started by removing the divider, and mice were allowed to make a decision (digging one ramekin) within 3 minutes. If no digging was performed within 3 minutes, the trial was scored as a null trial. Once mice started digging, the other ramekin was immediately removed from the behavioral box. If mice dug the correct ramekin to retrieve the reward (correct trial), a new trial would start once the reward was consumed. If mice dug the wrong ramekin embedded with the non-retrievable reward (incorrect trial), they would have a 1-minute timeout and a new trial would start.

A CCD camera (Basler acA1300-200um) was set above the behavioral box to capture the top-down view of mouse movements at 10 or 20 Hz, controlled by Pylon software. Video and electrophysiology recordings were synchronized via a common TTL pulse train (Arduino). Behavioral annotations were done manually post hoc.

Electrophysiology recordings were acquired at 20 kHz and hardware-filtered between 0.1-10 kHz (Intan Technologies). Signals were bandpass filtered between 300-6000 Hz and spikes were detected using a threshold of 4-8 standard deviations. The timestamp of the peak of each detected spike, as well as a 1.6 ms waveform centered at the peak, was extracted from each channel for offline spike sorting using MClust (Redish, 2014). Putatively duplicated units (peak correlation coefficient > 0.5 and 0 ms peak lag between spike rasters) were removed from further analysis. A recording session typically yielded 6-15 single units. A total of 161 single units were included in the analyses (inter-cluster distances > 20, cluster quality measure Lratio < 0.05). Cell type classification was based on trough to peak, full width at half maximum (FWHM) and baseline firing rate. Specifically, putative regular-spiking pyramidal neurons are identified by trough to peak > 0.5 ms and baseline firing rate < 10 Hz. Putative fast-spiking interneurons are identified by trough to peak < 0.5 ms and baseline firing rate > 10 Hz. The remaining units are considered unidentified.

In order to classify neuronal representations of different task-related variables, we performed Receiver-Operating-Characteristic (ROC) analysis on the firing rate of each unit for stimulus dimension, previous trial outcome and current trial choice separately. Dimension representation was defined as significant spiking responses between the odor-relevant stage (EDS) and combined digging medium-relevant stages (CD, REV and IDS) during ITI (−5 to 0 s from trial start) of the last 6 correct trials; a neuron was labeled ‘context+’ with the area under curve (AUC) > 0.5 and p < 0.05, conversely ‘context-’ neuron was defined with AUC < 0.5 and p < 0.05. Similar analysis was performed to classify outcome encoding in individual task stages, comparing spiking activity during ITI following correct trials against following incorrect trials. Removing the last 4 correct trials to better balance the number of correct and incorrect trials did not affect this analysis (data not shown). Choice classification was performed during a time window immediately prior to digging (−2 to 0 s from digging), comparing spiking activity preceding correct choices against preceding incorrect choices.

In order to classify neuronal representations of different task-related variables, we performed Receiver-Operating-Characteristic (ROC) analysis on the firing rate of each unit for stimulus dimension, previous trial outcome and current trial choice separately. Context representation was defined as significant spiking responses between the odor-relevant stage (EDS) and combined digging medium-relevant stages (CD, REV and IDS) during ITI (−5 to 0 s from trial start) of the last 6 correct trials; a neuron was labeled ‘context+’ with the area under curve (AUC) > 0.5 and p < 0.05, conversely ‘context-’ neuron was defined with AUC < 0.5 and p < 0.05. Similar analysis was performed to classify outcome encoding in individual task stages, comparing spiking activity during ITI following correct trials against following incorrect trials. Outcome encoding analysis was robust by removing the last 4 correct trials to better balance the number of correct and incorrect. Choice classification was performed during a time window immediately prior to digging (−2 to 0 s from digging) on each trial, comparing spiking activity preceding correct choices against preceding incorrect choices.

To assess the impact of different task-related variables on neuronal activity, a multilinear regression analysis was performed on the firing rate of each neuron (MATLAB function ‘fitglm’). Categorical regressors were context (odor - 1, digging medium - 0), outcome of previous trial (previous correct - 1, previous incorrect - 0), and choice of current trial (correct - 1, incorrect - 0). In Fig. S3, all trials (including incorrect trials) in CD, REV, IDS and EDS were pooled to estimate the coefficients. Model performance (fraction of variance explained, R^2^) of the complete model and the null model was compared using a permutation test: R^2^ values from the complete and null models were pooled, and then randomly assigned to two groups. The reported P values represented the proportion of iterations where the mean R^2^ difference between the two permutated groups exceeded the observed difference from 1000 iterations. Complete model R^2^ vs. null model R2, for Fig. S3 context+ neurons: 0.13 ± 0.028 vs. -0.0020 ± 0.0025, p < 0.001; context-neurons: 0.14 ± 0.021 vs. -9.6e-4 ± 0.0028, p < 0.001. In Fig. 5, 6 and Fig. S7-9, we estimated the coefficients of context, outcome, and choice by training and testing our model on data from CD, REV, IDS, and EDS stages. To estimate context coefficients, we pooled 80% of the trials (including incorrect trials) from all four stages for training and used the remaining 20% to test the model’s predictive performance on firing rates. Similarly, for estimating outcome and choice coefficients, we used 80% of the trials from each individual stage for training and the remaining 20% for testing. The models were evaluated using 5-fold cross-validation. To assess the model’s performance in predicting neuronal firing rates, we calculated the root mean square error (RMSE) for each temporal window. The RMSE values for choice and outcome in Fig. 5 and S7-8 are as follows: T1:1.26±0.13; T2:1.38±0.13; T3:1.63±0.19; T4:1.64±0.19. For context: T1:0.89±0.1; T2:0.91±0.11; T3:1.01± 0.1; T4:1.01±0.13. The RMSE values for choice and outcome in Fig. 6 and S9: T1:2.15±0.24; T2:1.87±0.21; T3:2.23±0.26; T4:1.92±0.26. Additionally, we calculated the Akaike Information Criterion (AIC) for the null model and compared it with the complete model. The comparison showed a significant difference between the complete model and the null model (complete model AIC: 59.45 ± 0.4 vs. null model AIC: 62.15 ± 0.44, p-value = 0.007). Similarly, for the context-specific model, there was a significant difference (context complete model AIC: 161.81 ± 2.04 vs. context null model AIC: 164.33 ± 2, p-value = 0.012).

For decoding analysis, we trained a linear multiclass error-correcting output codes (ECOC) model using support vector machine (SVM) binary learner and one-versus-one coding design (MATLAB function ‘fitcecoc’). We then used the MATLAB function ‘predict’ to examine decoding accuracy. For context decoding (Fig. 2), we used the last six correct trials in each stage (CD to EDS) to assess model prediction. For outcome and choice decoding (Fig. 3, 4), we used all trials in EDS to assess model prediction. Decoding analysis was performed using subsets of neurons (i.e., context-encoding, outcome-encoding, etc.) from individual recordings and comparisons were made between each recording and shuffled model. Due to relatively small number of trials in this task (c.f. Fig. 1B), we did not split the dataset into a training set and a testing set to examine decoding capacity. Instead, we shuffled class labels to establish chance level decoding accuracy. We note that chance level decoding probability may not be at 50%, as the shuffled model typically generated a prediction of uniform 0 or 1 states for all trials.

All data were presented as mean ± s.e.m. unless otherwise noted. Statistical tests were two-tailed signed rank for paired comparisons, and repeated-measure ANOVA for multiple comparisons unless otherwise noted.

