## Supplementary figures for "Distinct roles of prefrontal cortex neurons in set shifting"

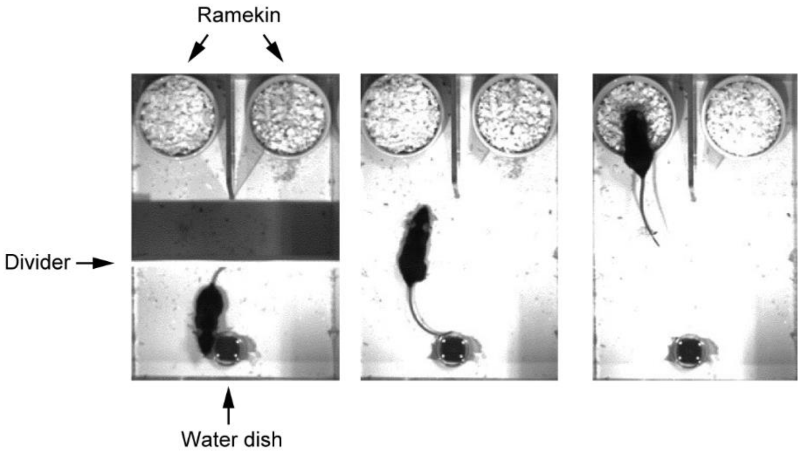


**Figure S1.**
Illustration of the behavioral box and three example video frames (left to right) showing moments when the mouse was in the waiting area (lower compartment) before trial start (divider inserted), approaching the ramekins after trial start (divider removed), and digging. The ramekin associated with the relevant stimulus was randomly placed in one of the two locations in each trial. This video was taken from a mouse without tetrode implant.

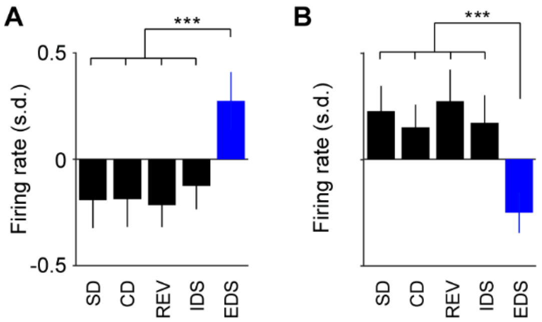


**Figure S2.**

(A) The context+ neurons identified in Fig. 2 exhibited similar responses in SD as in other digging medium-relevant stages. Repeated-measures ANOVA, F(4, 104) = 13.6, P = 5.6e-9, n = 27. Post hoc Tukey-Kramer tests: EDS vs. SD, P = 2.7e-6; EDS vs. CD, P = 3.1e-4; EDS vs. REV, P = 5.6e-6; EDS vs. IDS, P = 3.7e-4. All other paired tests were not significant.

(B) The context- neurons identified in Fig. 2 exhibited similar responses in SD as in other digging medium-relevant stages. Repeated-measures ANOVA, F(4, 88) = 11.2, P = 2.1e-7, n = 23. Post hoc Tukey-Kramer tests: EDS vs. SD, P = 5.1e-5; EDS vs. CD, P = 1.3e-5; EDS vs. REV, P = 1.1e-5; EDS vs. IDS, P = 7.7e-5. All other paired tests were not significant.

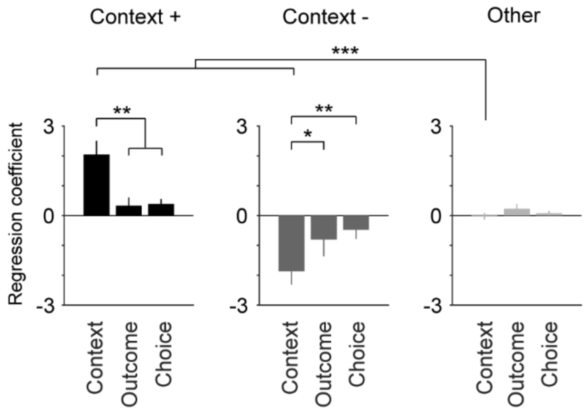


**Figure S3.**

The regression coefficients of stimulus dimension (context) in context neurons were stronger than the coefficients of other variables (outcome and choice), and stronger than those of non-context neurons. Context+ neurons: Repeated-measures ANOVA, $F(2, 52) = 8.96$, $P = 4.5e-4$, $n = 27$. Post hoc Tukey-Kramer tests: Context vs. Outcome, $P = 0.01$; Context vs. Choice, $P = 0.0082$; Outcome vs. Choice, $P = 0.98$.

Context- neurons: Repeated-measures ANOVA, $F(2, 44) = 6.2$, $P = 0.0042$, $n = 23$. Post hoc Tukey-Kramer tests: Context vs. Outcome, $P = 0.018$; Context vs. Choice, $P = 0.0027$; Outcome vs. Choice, $P = 0.79$.

The coefficients of stimulus dimension: Context+ vs. Other, $P < 0.001$; Context- vs. Other, $P < 0.001$. Permutation test.

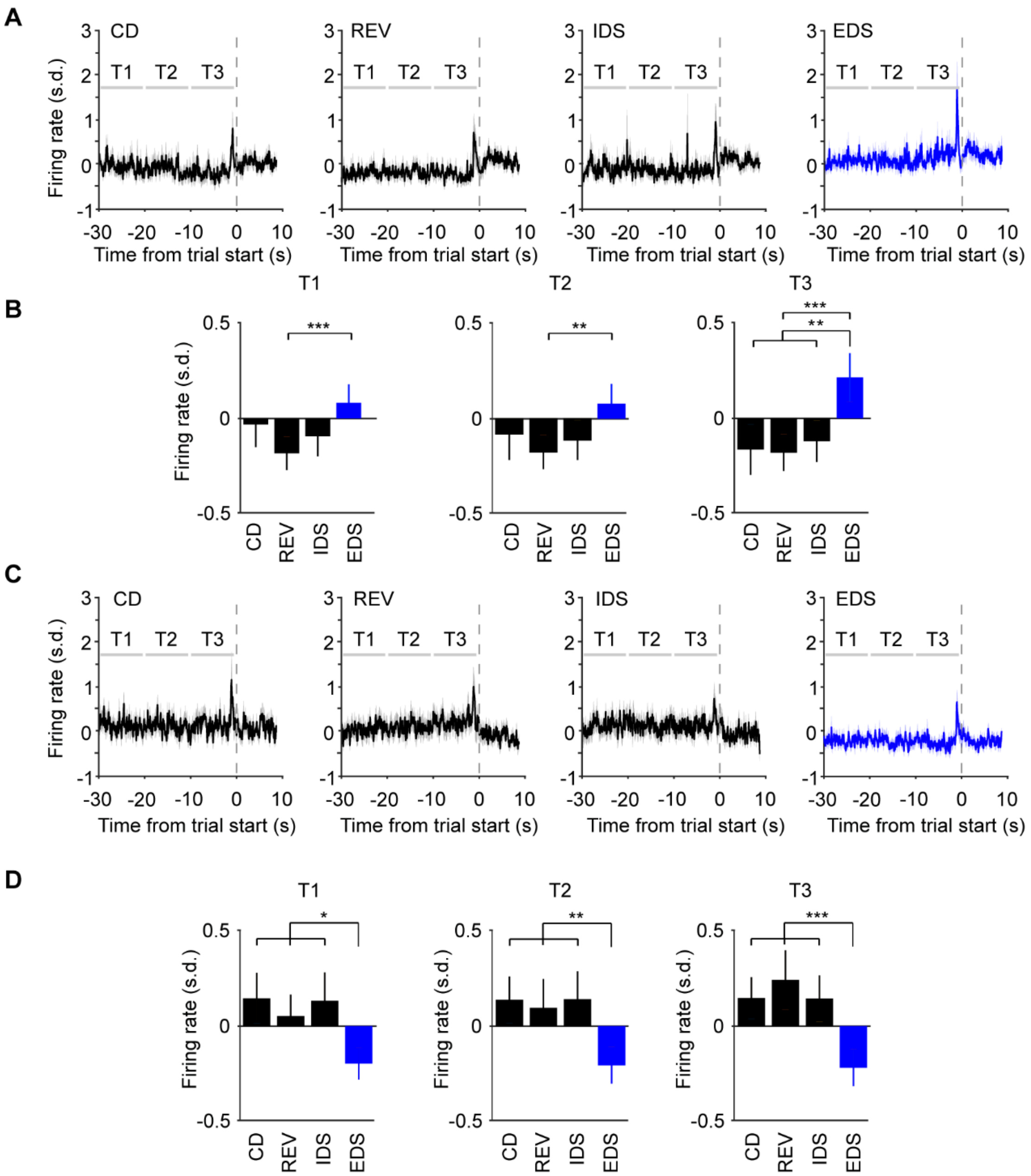


**Figure S4.**
(A) Group mean peri-event spike time histogram (PETH) of context+ neurons (n = 27) aligned to trial onset in stages CD through EDS. Mean firing rate quantified in 3 10-s time windows before trial start (grey horizontal bars. T1: -30 to -20 s; T2: -20 to -10 s; T3: -10 to 0 s from trials start) is shown in B.
(B) T1: Repeated-measures ANOVA, $F(3, 78) = 4.6$, $P = 0.005$, n = 27. Post hoc Tukey-Kramer tests: EDS vs. REV, P = 7.04e-5. All other paired tests were not significant. T2: Repeated-measures ANOVA, $F(3, 78) = 3.5$, $P = 0.018$, n = 27. Post hoc Tukey-Kramer tests: EDS vs. REV, P = 0.005. All other paired tests were not significant. T3: Repeated-measures ANOVA, $F(3, 78) = 12.7$, P = 8.02e-7, n = 27. Post hoc Tukey-Kramer tests: EDS vs. CD, P =0.001; EDS vs. REV, P = 8.6e-6; EDS vs. IDS, P = 0.001. All other paired tests were not significant.

(C) Group mean peri-event spike time histogram (PETH) of context- neurons (n = 23) aligned to trial onset in stages CD through EDS. Mean firing rate quantified in 3 different time windows before trial start (grey horizontal bars. T1: -30 to -20 s; T2: -20 to -10 s; T3: -10 to 0 s from trials start) is shown in D.
(D) T1: Repeated-measures ANOVA, $F(3, 66) = 4.9$, $P = 0.0039$, n = 23. Post hoc Tukey-Kramer tests: EDS vs. CD, P = 0.046; EDS vs. REV, P = 0.015; EDS vs. IDS, P = 0.015. All other paired tests were not significant. T2: Repeated-measures ANOVA, $F(3, 66) = 8.5$, $P = 7.3e-5$, n = 23. Post hoc Tukey-Kramer tests: EDS vs. CD, P = 0.0004; EDS vs. REV, P = 0.0055; EDS vs. IDS, P = 0.0013. All other paired tests were not significant. T3: Repeated-measures ANOVA, $F(3, 66) = 11.07$, $P = 5.6e-6$, n = 23. Post hoc Tukey-Kramer tests: EDS vs. CD, P = 6.12e-5; EDS vs. REV, P = 1.2e-4; EDS vs. IDS, P = 4.4e-4. All other paired tests were not significant.

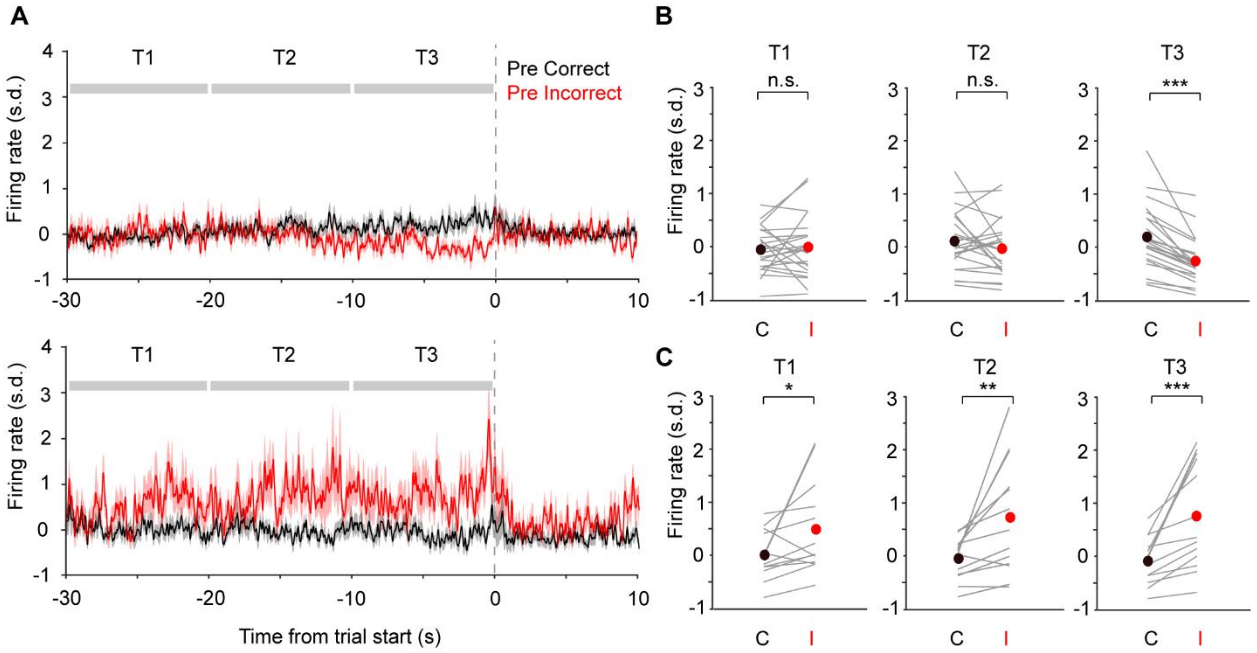


**Figure S5.**
(A) Group mean peri-event spike time histogram (PETH) of outcome+ neurons in EDS (n = 23) aligned to trial onset when previous trials were correct (black) and incorrect (red). Mean firing rate quantified in 3 10-s time windows before trial start (grey horizontal bars. T1: -30 to -20 s; T2: -20 to -10 s; T3: -10 to 0 s from trials start) is shown in B.
(B) The activity of outcome+ neurons in the intertrial interval following correct (black) and incorrect (red) trials differed in the last 10 seconds before trial onset (T1: P =0.60; T2: P =0.26; T3: P = 2.7e-5, n = 23).
(C) Group mean peri-event spike time histogram (PETH) of outcome- neurons in EDS (n = 13) aligned to trial onset when previous trials were correct (black) and incorrect (red). Mean firing rate quantified in 3 10-s time windows before trial start (grey horizontal bars. T1: -30 to -20 s; T2: -20 to -10 s; T3: -10 to 0 s from trials start) is shown in D.
(D) The activity of outcome- neurons in the intertrial interval following correct (black) and incorrect (red) trials differed in the last 20 seconds before trial onset (T1: P =0.02; T2: P =0.002; T3: P = 2.4e-4, n = 13).

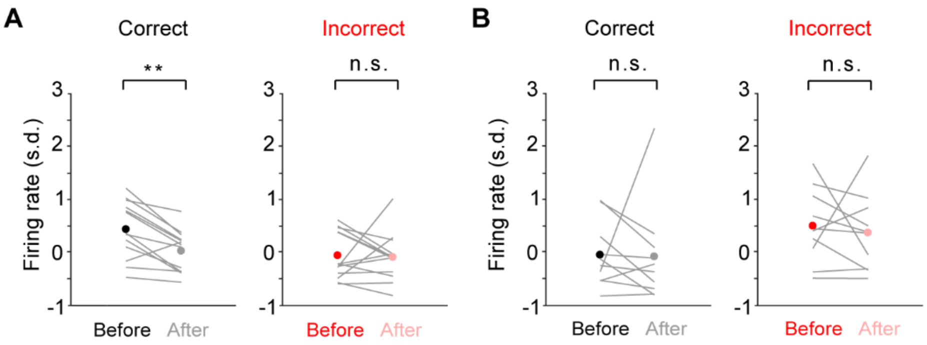


**Figure S6.**
(A) The activity of choice+ neurons in EDS (n = 13) before and after digging (-2 to 0 s vs. 0 to 2 s from digging) only differed in correct trial (Left, P = 0.002) but not incorrect trials (Right, P = 0.5).
(B) The activity of choice- neurons in EDS (n = 10) before and after digging (-2 to 0 s vs. 0 to 2 s from digging) did not differ in correct trial (Left, P = 0.28) or incorrect trials (Right, P = 0.38).

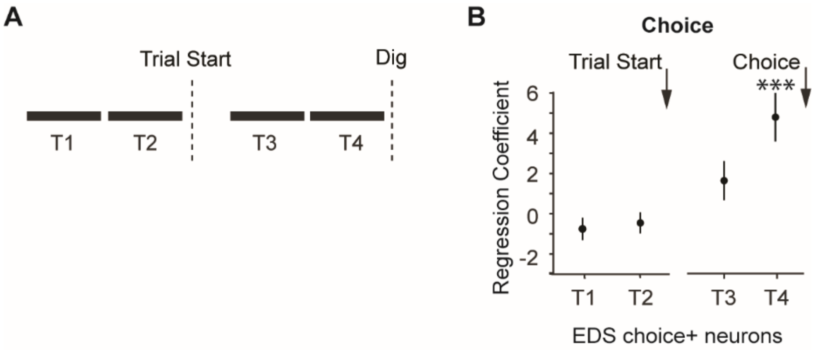


**Figure S7.**
(A) Illustration of the four trial epochs. T1: -4 to -2 s from trial onset; T2: -2 to 0 s from trial onset; T3: -4 to -2 s from digging, T4: -2 to 0 s from digging;
(B) Regression coefficients of the choice factor for choice + neurons in EDS (n = 13). Coefficients in T4 were significantly different from 0. T1, p = 0.15; T2, p = 0.31; T3, p = 0.13; T4, p = 2e-4.

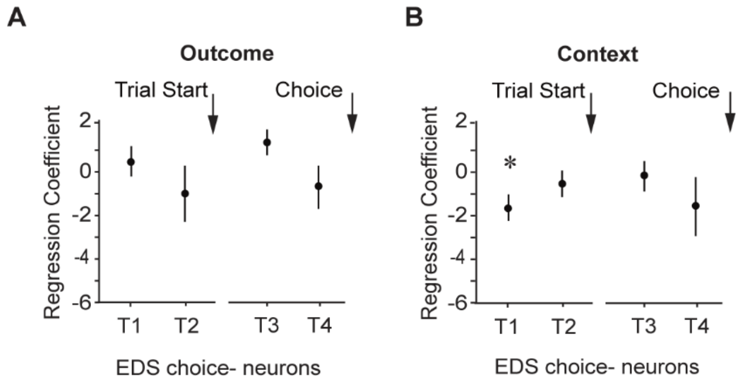


**Figure S8**

(A) Regression coefficients of the outcome factor for choice- neurons in EDS (n = 10). Coefficients in all windows were not different from 0. T1, P = 0.58; T2, P = 0.41; T3, P = 0.07; T4, P = 0.5.

(B) Regression coefficients of the context factor for choice- neurons in EDS (n = 10). Coefficients in T1 were different from 0. T1, P = 0.023; T2, P = 0.28; T3, P = 0.61; T4, P = 0.23.

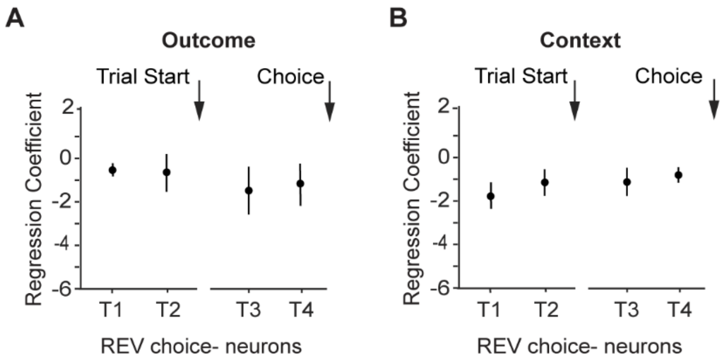


**Figure S9**

(A) Regression coefficients of the outcome factor for choice- neurons in REV (n = 6). Coefficients in all windows were not different from 0. T1, P = 0.65; T2, P = 0.94; T3, P = 0.45; T4, P = 0.57.

(B) Regression coefficients of the context factor for choice- neurons in REV (n = 6). Coefficients in all windows were not different from 0. T1, P = 0.1; T2, P = 0.47; T3, P =0.6; T4, P = 0.79.
